# Ex vivo and in vivo imaging of mouse parietal association cortex activity in episodes of cued fear memory formation and retrieval

**DOI:** 10.1101/863589

**Authors:** Olga I. Ivashkina, Anna M. Gruzdeva, Marina A. Roshchina, Ksenia A. Toropova, Konstantin V. Anokhin

## Abstract

The parietal cortex in rodents has an integrative function and participates in sensory and spatial processing, movement planning and decision-making. However, much less is known about its functions in associative memory processing. Here using Fos immunohistochemical mapping of neuronal activity and two-photon imaging in Fos-eGFP mice we show an involvement of anterior part of the parietal cortex (PtA) in the formation and retrieval of recent fear memory in mice. Using ex vivo c-fos imaging we demonstrate the specific activation of the PtA during recent memory retrieval. In vivo two-photon c-fos imaging confirms these results as well as establishes the activation of the PtA neurons during fear memory formation. Additionally, we describe a design of Fos-Cre-GCaMP transgenic mice to investigate long-term changes of calcium dynamics in neurons captured with Fos-TRAP technique during fear conditioning training.

## Introduction

The parietal cortex is an association cortical area that has been involved in multisensory processing, navigation, motion planning and decision-making. Though this area has been well studied in behavioral tasks in primates (Lynch, 1980), only more recently it has become the subject of corresponding analysis in rodents (Lyamzin, Benucci, 2019, Hovde et. al, 2018).

Anatomically the rodent parietal cortex has connections with diverse brain areas, in particular with other association cortical regions, such as orbitofrontal, retrosplenial and anterior cingulate cortices (Zingg et al., 2014, Hovde et. al, 2018). Its neurons respond to complex or modality-specific (auditory, visual or somatosensory) stimuli (Lippert et al., 2013, Mohajerani et al., 2013). In the mouse brain this region is defined as an area between visual and somatosensory cortices, and consists of anterior and posterior parts (Franklin, Paxinos, 1997, Dong, 2008, Lyamzin, Benucci, 2019). According to Franklin and Paxinos Mouse Brain Atlas (1997), the anterior part of parietal cortex corresponds to parietal association cortex (PtA). Network analysis of cortical areas connectivity revealed that the PtA has high level of betweenness centrality that indicated PtA as a strong hub region within the network (Lim et al., 2012). However, most of the existing studies address the posterior parietal cortex but not its anterior associative part. In this study we address specifically the PtA.

Functionally the rodent parietal cortex is involved in space navigation (Whitlock et. al, 2014) and decision-making processes (Goard et al., 2016, Zhong et al., 2019). Other association cortical areas, which send projections to the PtA, are involved in coding and retrieval of the different types of memory (retrosplenial (Vann et al., 2009, Czajkowski et al., 2014), cingulate (Goshen et al., 2011, Frankland et al., 2004) and frontal associative (Nakayama et al., 2015). The PtA also participates in the retrieval of recent and remote spatial memory (Save, Moghaddam, 1996, Maviel et al., 2004). However, the contribution of this cortical area to associative memory is still poorly understood. In this study we address this question using c-Fos neuronal activity mapping, two-photon microscopy and calcium imaging.

Expression of immediate early genes (IEGs), such as *c-fos* is commonly used to identify neurons which were activated by learning and are specifically involved in memory encoding (Josselyn et al., 2015, Tonegawa et al., 2015). *C-fos* encodes the transcription factor that regulates activity of effector genes and the following long-term plasticity in neurons. Different Fos-based methods are used to investigate the experience-induced changes of brain neuronal circuits during various memory processed. However, c-Fos immunostaining can be used to access the neuronal activity only at a single timepoint. In contrast, in vivo Fos-imaging approaches allow observing the involvement of the same brain area in different behavioral episodes. In this case, Fos-GFP transgenic mice can be used for repeated imaging of Fos-positive neurons and comparing neuronal populations which were active during learning or memory retrieval (Barth et al., 2004, Czajkowski et al., 2014, Milczarek et al., 2018). Besides, the method of targeted recombination in active populations (TRAP) can be used to capture the candidate engram neurons (Guzowski et al., 1999, Josselyn et al., 2015). TRAP is an approach to obtain permanent genetic access to distributed neuronal ensembles that are activated by experiences within a limited time window (Guenthner et al., 2013, Ivashkina et al., 2018). This method uses mice in which the tamoxifen-dependent recombinase Cre^ERT2^ is expressed in an activity-dependent manner under the Fos promotor control. Active cells that express Cre^ERT2^ can undergo recombination only in presence of tamoxifen (TM), allowing genetic access to neurons that were active during a time window of less than 12 h. Cre^ERT2^ is retained in the cytoplasm of active cells and recombination cannot occur in the absence of TM. Nonactive cells do not express Cre^ERT2^ and do not undergo recombination, even if TM was presented (Guenthner et al., 2013). Previously TRAP approach was used to assess impact of neurons, genetically captured during context fear conditioning on subsequent memory retrieval (Denny et al., 2014). Optogenetic silencing of functional labeled neuronal populations in CA3 or DG prevented expression of the corresponding memory. In the present study we use these approaches for investigation of PtA cortex involvement in associative fear memory formation and recent retrieval.

## Materials and methods

### 1.1. Animals

Male C57Bl/6J mice (2–3 months old) were used for ex vivo study of c-Fos expression in the PtA after fear conditioning (FC) or memory retrieval. Transgenic Fos-EGFP male and female mice (B6.Cg-Tg(Fos/EGFP)1-3Brth/J, JAX Stock No: 014135 The Jackson Laboratory) were used for in vivo two-photon investigation of specific activation of PtA neurons during FC or memory retrieval. Transgenic Fos-Cre-GCaMP male and female mice were obtained by crossing two transgenic mouse lines Ai38 (RCL-GCaMP3) (B6;129S-*Gt(ROSA)26Sor*^*tm38(CAG-GCaMP3)Hze*^/J, JAX Stock No: 014538 The Jackson Laboratory) and Fos^CreER^ (B6.129(Cg)-*Fos*^*tm1.1(cre/ERT2)Luo*^/J, JAX Stock No: 021882 The Jackson Laboratory).

Wild type mice were group housed 5-6 per cage. Transgenic mice were housed individually. All animals were kept under a 12 h light/dark cycle. All experiments were performed during the light phase of the cycle. All methods for animal care and all experiments were approved by the National Research Center “Kurchatov Institute” Committee on Animal Care (protocol No. 1, 7 September 2015) and were in accordance with the Russian Federation Order Requirements N 267 M3 and the National Institutes of Health Guide for the Care and Use of Laboratory Animals.

### 1.2. Behavior

During sound fear conditioning mice were placed into the fear conditioning chamber (Med Associates) for 3-minute exploration of context A, then they were presented with seven sound signals and each signal was followed by a foot shock (2 sec, 0.75 mA (for wild type mice) or 1 mA (for the transgenic mice)). The sound signal contained 5 tones (2 sec, 9 kHz, 80 dB) with 2 sec intervals and different interstimulus intervals (40 s or 60 s) between these seven signals. 24 h later cue memory was tested in context B. Mice were placed in the FC chamber for 3-minute exploration, and then they were presented with the conditioned stimulus (CS) for 3 minutes, which consisted of 45 sound signals (2 sec, 9 kHz, 80 dB) with 2 sec intervals. Context A and B were cleaned before and after each session with 70% ethanol or 53% ethanol solution of pepper mint, respectively. Context A consisted of a plastic box (30 cm × 23 cm × 21 cm) with metal rods on the floor, chamber white light was off, with white light in the room. Context B was different from the training context and had a plastic sheet with wood sawdust on the floor and black plastic walls placed at 60° to the floor, forming a triangular roof. Moreover, in Context B the white chamber’s light was on, and red light illuminated the room. Freezing behavior was quantified using an automatic detection system (Video fear conditioning system (MED Associates Inc.)) and Video Freeze (MED Associates Inc.).

Wild type mice were divided into the three groups: Shock (N(FC) = 12, N(test) = 11), No shock (N(FC) = 10, N(test) = 11) and Home cage group (N(FC) = 12, N(test) = 13). Mice from the Shock group were fear conditioned using the protocol described above. Mice from the No shock group were submitted to the same procedure that was for the Shock group without foot shock. Half of the trained mice were decapitated 90 minutes after FC (N(FC)), while the half was tested and decapitated 90 minutes after memory test (N(test)). The Home cage group mice were sacrificed at the same time.

Fos-EGFP mice were divided into the same groups (N(Shock) = 7, N(no shock) = 6, N(home cage) = 3). Mice from the Shock group were submitted to the auditory FC one month after cranial window surgery. 24 h after the same mice were tested in the new context B. We imaged the same area of PtA three days before FC, 90 minutes after FC and 90 minutes after memory test. No shock group mice were submitted to the same procedure without foot shock. The Home cage group mice were taken to imaging from the home cage without any prior procedure.

For Fos-Cre-GCaMP mice FC was performed one day after TM injection in context A with the described protocol.

### 1.3. Genotyping

Fos-Cre-GCaMP mice were genotyped at 30-60 days age. For genotyping we extracted DNA from the tail tissue in lysis buffer (0.01 M Tris-HCl, pH=7.5; 0.01 M EDTA, pH=8.0; 0.1 M NaCl; 1% SDS; 0.2 mg/ml proteinase K). We performed PCR (in PCR buffer with Taq-polymerase, DNTP, forward and reverse primers) and identified DNA sites using electrophoresis in agarose gel (2.5%). We used primers oIMR4981, oIMR8038, 34319, 34962 for GCaMP genotyping, and 17016-17018 for Fos-Cre genotyping (The Jackson Laboratory).

### 1.4. Surgery

For cranial window surgery transgenic mice were anesthetized with an intraperitoneal injection of zoletil (0.04 mg/g body weight) and xylazine (0.5 µg/g body weight). Dexamethasone (4 mg/kg) was administered subcutaneously five minutes before surgery to prevent tissue stress and cerebral edema. Viscotears (Novartis Healthcare) moisturizing gel was applied to prevent eye drying. Mice were fixed in stereotaxic frame (Stoelting) and 37°C body temperature was maintained by a heating plate (Physitemp). The parietal association cortex (PtA) was determined in accordance with coordinates from the anatomical mouse brain atlas (centered 1.0 mm lateral and 1.7 mm posterior to the Bregma). Craniotomy (3 mm diameter) over the PtA was performed in accordance with a previously described protocol 14.. A 5-mm round glass coverslip (Menzel, Thermo Fisher) was attached to the skull using cyanoacrylate glue for glass (Henkel). A Neurotar (Neurotar Ltd.) head post was cemented to the skull with dental cement (Stoelting) and was later used for fixation in the Mobile Home Cage (MHC).

Two weeks after surgery Fos-EGFP and Fos-Cre-GCaMP mice were head-fixed each day starting from 5 minutes to 40 minutes during two weeks for habituation to conditions of imaging. For fixation Fos-Cre-GCaMP mice the Mobile Home Cage system (MHC) was used. The MHC consisted of a carbon fiber cage (18.5 cm in diameter, 7 cm height walls) that floated on an air-flows-above the surface of an air dispenser, and with pillars and a bridge for head fixation (Neurotar, Ltd.).

### 1.5. Tamoxifen induction in the Fos-Cre-GCaMP mice

The Fos-Cre-GCaMP mice received single tamoxifen (TM) (Sigma) injection intraperitoneally (150 mg/kg) one day before training. Tamoxifen was dissolved in the corn oil (Sigma) (10 ml/kg) and 96% ethanol (1,3 ml/kg) by shaking at 65°C for 1-2 h. This dose and timing of injection was used in accordance with previously described protocol (Holtmaat et al., 2009).

### 1.6. Immunohistochemistry

Brains were extracted and immediately frozen in liquid nitrogen vapor. Brains were sectioned into 20 um thick coronal sections using a cryostat (Leica), fixed in 4% paraformaldehyde and then stained for c-Fos. For this purpose, we used primary rabbit polyclonal antibodies against c-Fos protein (sc-52, Santa Cruz Biotechnology, dilution 1:500) and horse secondary antibodies against rabbit conjugated with avidin-biotin complex (ImPRESS reagent kit anti-rabbit, Vector Laboratories) in 0.06% diaminobenzidine solution (DAB, Sigma). Imaging of tissue was performed with a fluorescence virtual slide microscopic scanner (Olympus, VS110).

### 1.7. *In vivo* two-photon imaging

Two-photon imaging of neurons in Fos-EGFP and Fos-Cre-GCaMP mice was performed 30–60 days after surgery using an Olympus MPE1000 two-photon microscope equipped with a Mai Tai Ti:Sapphire femtosecond-pulse laser (Spectra-Physics) and a water-immersion objective lens with 1.05 NA (Olympus). 960 nm wavelength was used for excitation. Series of images were recorded with the Olympus Fluoview Software Version 3.1. Fos-Cre-GCaMP mice were placed in the MHC.

#### Fos-EGFP mice

For the Fos-EGFP mice the volume series of images (0-350 μm under the pia) were recorded at 0.82 frames per second continuously, with a field of view of 500×500 μm with 512×512 pixels. We performed three sessions of two-photon imaging of the PtA. The first one was performed three days before learning when the mice were taken from the home cages. The second and the third sessions were performed 90 minutes after FC or memory test respectively. The same area of visualization to repetitive imaging on different days was found by following the blood vessels on the brain surface.

#### Fos-Cre-GCaMP mice

To determine the dynamics of GCaMP accumulation after the Cre recombination, we visualized the PtA (0-300 μm under the pia) at different time points after the FC training in the Fos-Cre-GCaMP mice.

Volume series of images (0-400 μm under the pia) were recorded 0.2 frames per second continuously, with a field of view of 317×317 μm with 512×512 pixels three days after FC before the calcium imaging of PtA L2/3 neurons (approximately 100-200 μm deep from pia). During the calcium imaging mice received seven series of the CS (one series consisted of 10 short tones (2 sec, 9 kHz, 80 dB) with 2 sec intervals). Time series of images were recorded at 1.78 frames per second continuously, with a field of view of 253×253 μm with 256×256 pixels and zoom 2x. For spontaneous calcium activity in dendrites we recorded a time series of images at 1.92 frames per second with a field of view of 35×44 μm with 180×144 pixels and zoom 4x at the depth of 5-10 μm without any stimulation.

The image analysis was performed using the Olympus Fluoview Software Version 3.1, FIJI, Imaris 7.4.2 and a custom Python plugin. For calcium transients analysis we estimated changes of fluorescence signal (ΔF/F). For calculation of the ΔF/F ratio, regions of interest (ROIs) were first drawn around individual neuronal somata or dendritic spines. ΔF/F were calculated by subtracting each value with the mean of the lower 50% of previous 10-s values and dividing it by the mean of the lower 50% of previous 10-s values (Miller et al., 2014).

## Results

### Ex vivo investigation of c-Fos expression in PtA

We first analyzed c-Fos expression in the PtA during fear memory formation and retrieval. Wild type mice were trained to associate seven acoustic stimuli (conditioned stimuli, CS) with foot shocks. Only the Shock group developed freezing behavior as the number of associative pairings increased (Fig. 1, A). Freezing response to the CS presentation during memory test (24 h after fear conditioning (FC)) was higher in the Shock group compared with No shock group (two-way ANOVA, p<0.0001) (Fig. 1, B). The freezing level during novel context exploration, however, was the same in the Shock and No shock groups (two-way ANOVA, p<0.0001), (Fig. 1, B). These results demonstrate that mice formed associative memory to the CS and did not exhibit generalization to a novel environment.

**Fig. 1.**
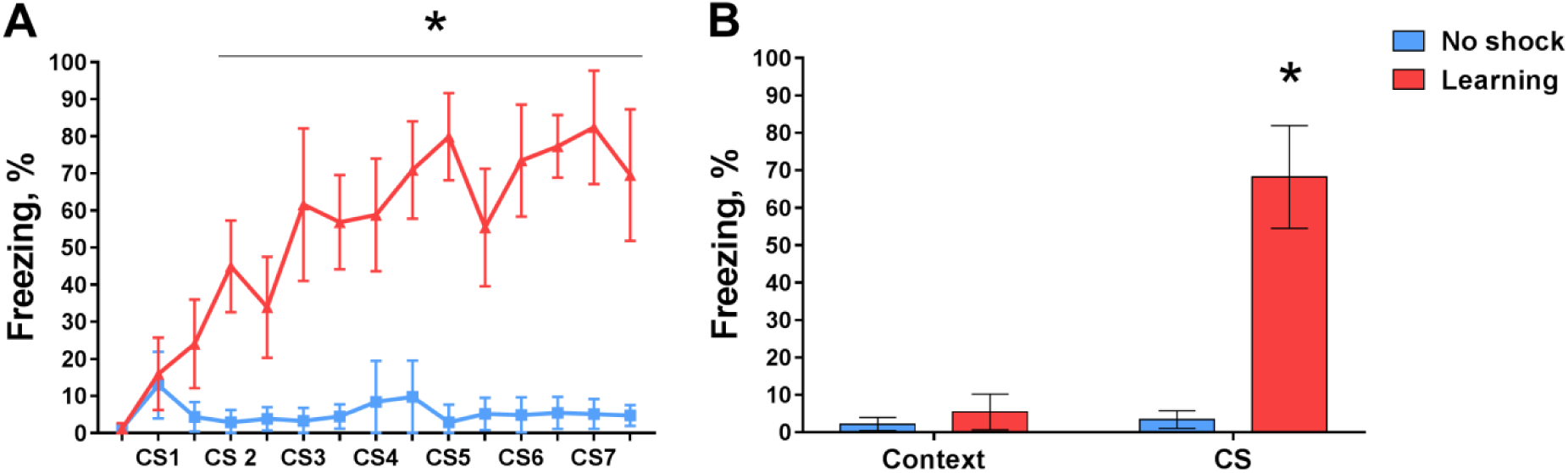
Freezing levels during fear learning (A) and memory retrieval (B) (Mean, 95% CI, n=11 per group). Freezing level is increasing only in the Learning group (*, two-way ANOVA, p<0.0001). Freezing response to presentation CS in test is higher in the Learning group compared with the No shock group or compared with freezing of the Learning group before CS presentation (*, two-way ANOVA, post hoc Tukey test, p<0.0001).

To investigate the activation of PtA during learning or memory retrieval we used c-Fos immunohistochemistry staining of wild type mice brain sections 90 minutes after FC or memory retrieval. We found that the number of c-Fos-positive cells in the PtA was higher in the Shock and No shock groups than in the HC group during FC and memory retrieval (one-way ANOVA, p<0.001) (Fig. 2, A). The number of active cells was the same in the Shock and No shock groups during FC. However, during memory test, the number of c-Fos-positive cells was higher in the Shock group comparing with the No shock group (one-way ANOVA, p=0.0364) (Fig. 2, B). This data suggest that PtA neurons are active during new experience acquisition and are specifically involved in the session of recentCS memory retrieval.

**Fig. 2.**
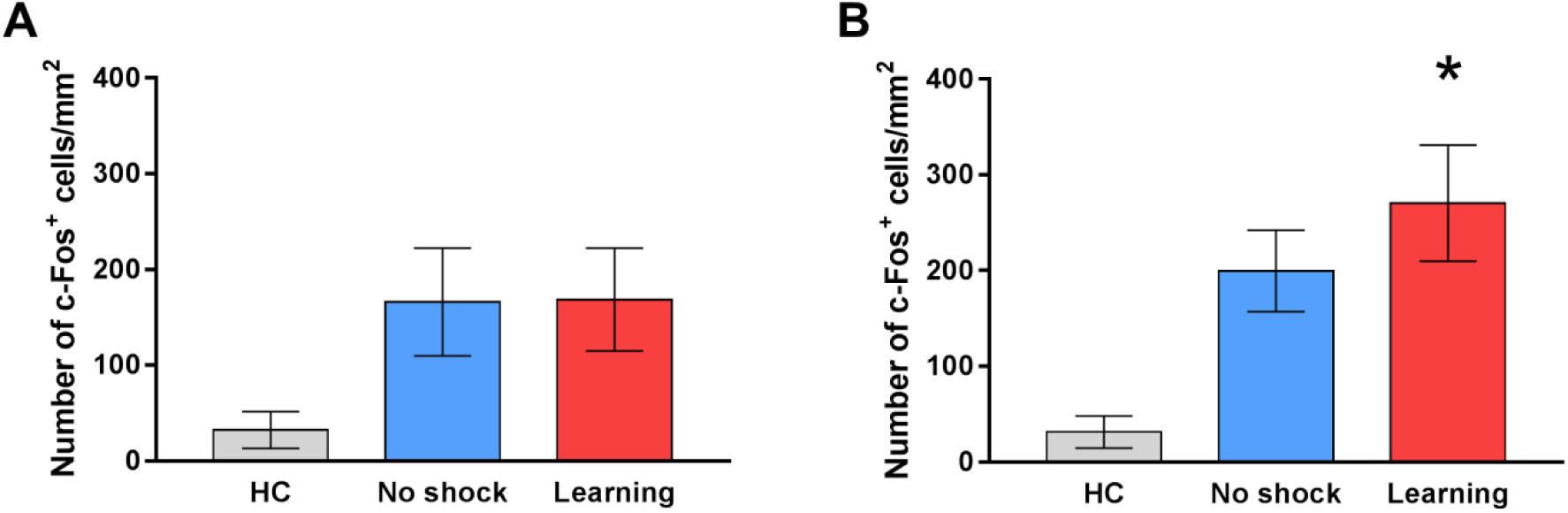
The number of c-Fos-positive cells in the PtA. A: Fear learning, B: Fear memory retrieval (Mean, 95% CI, n=10-13 per group). The number of c-Fos-positive cells in the PtA is higher in the Learning and the No shock groups than in the HC group during learning and memory retrieval (one-way ANOVA, post hoc Tukey test, p<0.001). During memory retrieval the number of c-Fos-positive cells is higher in the Learning group compared with the No shock group (*, one-way ANOVA, post hoc Tukey test, p=0.0364).

### In vivo investigation of c-Fos expression in the PtA of Fos-EGFP mice

Next, we determined activation of the same PtA neurons in FC and memory retrieval. For this purpose, we performed in vivo two-photon imaging of the same PtA area before FC, after FC and after memory test in Fos-eGFP transgenic mice. This approach allowed us to address the specific neuronal population after different behavioral procedures.

Overall, we identified 9325 neurons in all the mice during all sessions of two-photon visualization. Out of the all identified neurons, 17% changed their activity (i.e. were activated or inactivated) at least during one imaging session. We found different number of Fos-EGFP positive neurons in 16 mice in control imaging session (before learning), resulting in the man number of 521.5±155.9 neurons per mouse (mean±95% CI).

To investigate the dynamics of the Fos-active neurons we compared the number of such activated neurons normalized to the number of neurons which were active during the first imaging session performed before the FC trial for each group. The number of Fos-active neurons increased during the FC and memory recall in the Shock group but not in the control groups (t-test, comparing with 1, p=0.0156) (Fig. 3).

**Fig. 3.**
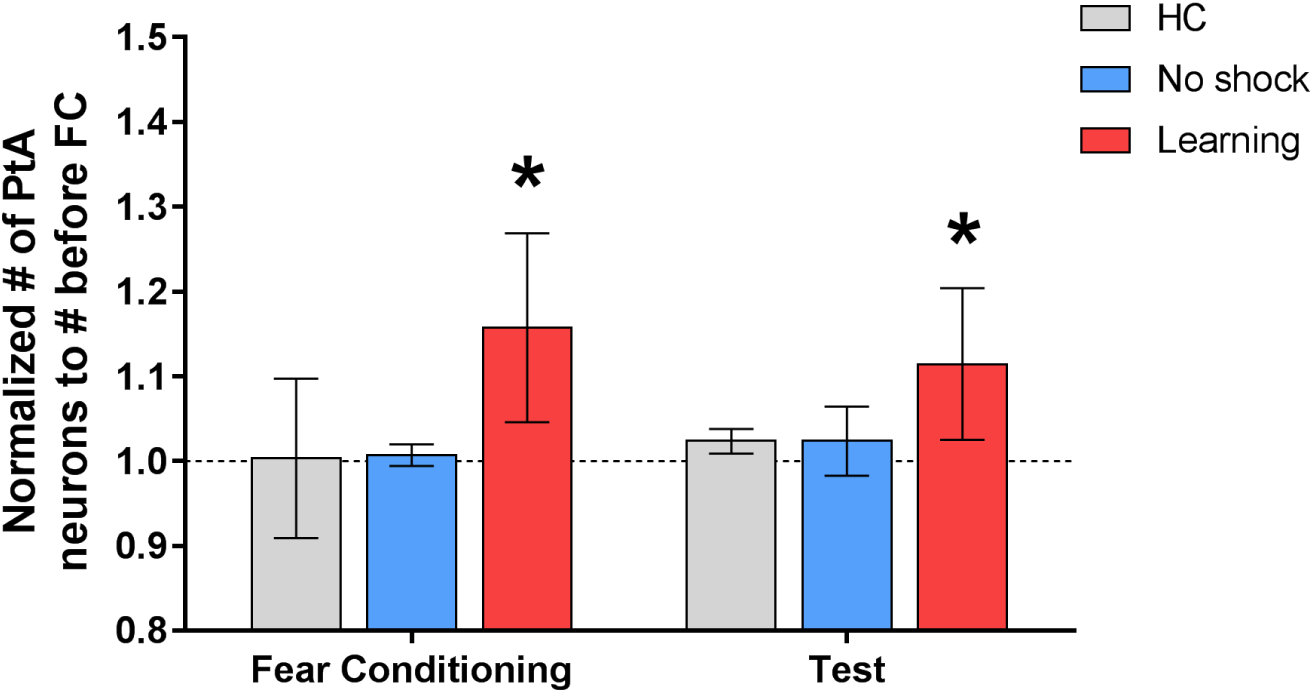
Number of the PtA neurons normalized to the number of neurons which were active during first visualization (before FC) (Mean, 95% CI, n(HC)=3, n(No shock)=6, n(Learning)=6). The number of active neurons was increased during learning and memory retrieval in the Learning group but not in the control groups (* - compared with 1, t-test, p=0.0156).

Next, we compared the number of neurons which were inactive before the FC but were active during the FC or memory test. The number of neurons which were activated during FC was higher in the Shock group than in the control groups (one-way ANOVA, p<0.05) (Fig. 4, A). The number of neurons that were inactive before FC but were activated by memory retrieval was similar in the all groups (Fig. 4, B).

**Fig. 4.**
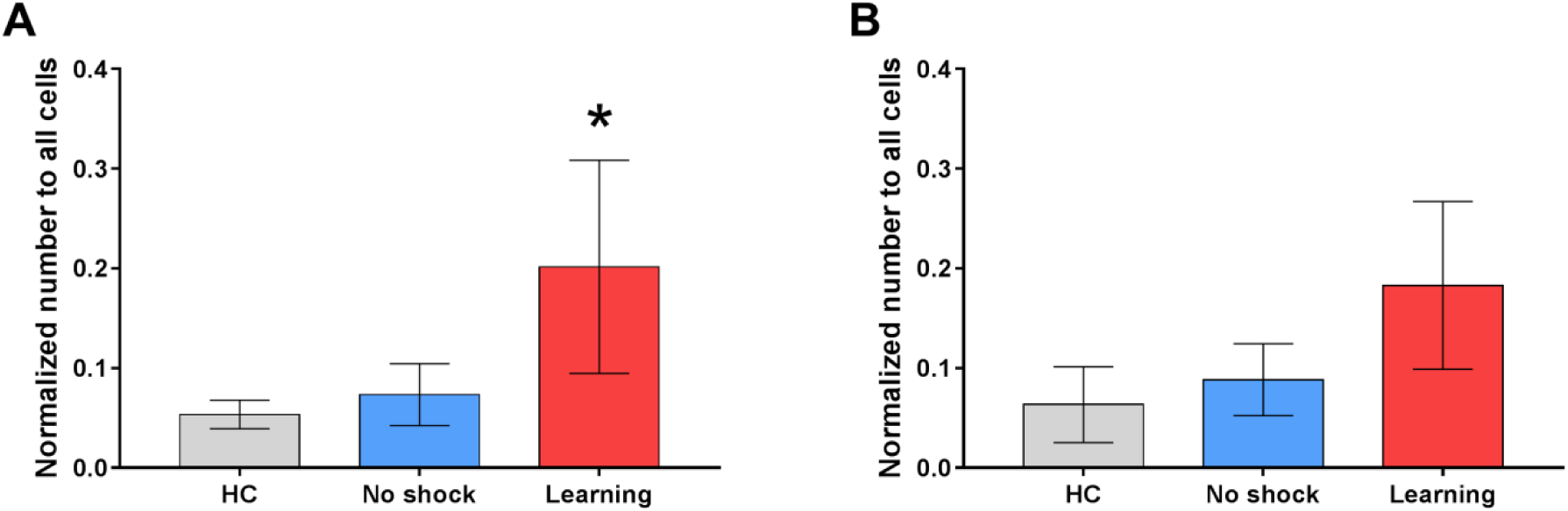
Number of the active PtA neurons, which were inactive before FC, normalized to the all identified neurons. A: Fear learning, B: Memory retrieval (Mean, 95% CI, n(HC)=3, n(No shock)=6, n(Learning)=6). The number of neurons which were inactive before FC and were active during FC was higher in the Learning group than in the control groups (*, one-way ANOVA, post hoc Tukey test, p<0.05).

Collectively, this result indicate that PtA neurons are involved in the sessions of formation and retrieval of associative fear memory.

To address this issue in a more precise and time-resolved manner, we recorded calcium activity specifically in neurons, which were involved in episodes of memory formation. For these purposes we crossed Fos^CreER^ and Ai38 (RCL-GCaMP3) transgenic mice to utilize the TRAP strategy to label Fos-positive neurons during FC learning with the GCaMP calcium sensor.

### Characterization of Fos-Cre-GCaMP transgenic mice

As was expected, we found that DNA of the Fos-Cre-GCaMP transgenic mice contain specific sites for GCaMP (226 bp) and Fos-Cre (293 bp) compared 128 bp and 215 bp from DNA of wild type mouse (Fig.5).

**Fig. 5.**
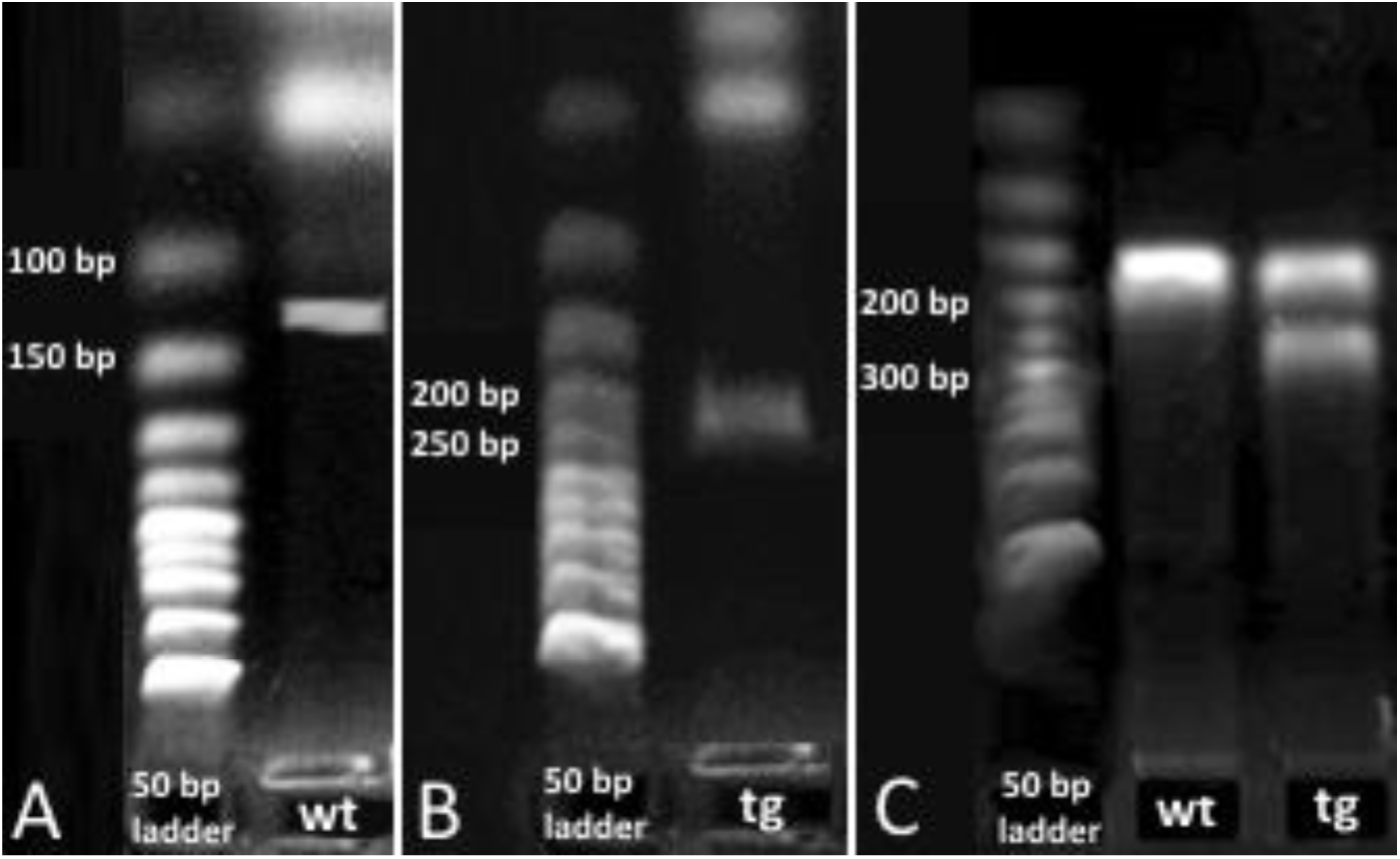
Electrophoresis of the Fos-Cre-GCaMP transgenic mouse DNA (tg) and DNA of the wild type mouse (wt) and 50 bp ladder. A, B: The specific site for gcamp (226 bp) compared 128 bp of for DNA wild type mouse. C: The specific site for fos-cre (293 bp) compared 215 bp for DNA of wild type mouse.

To determine time course of GCaMP accumulation in TRAPed neurons we performed repeated two-photon imaging of the PtA area at different time points after the session of FC-triggered Cre-recombination. The first visualization was performed two hours after FC to investigate the possibility of the early spontaneous recombination. We found no GCaMP-expressing neurons in PtA. Thus, no background or spontaneous recombination events appear in short time after tamoxifen injection. One day after FC the number of neurons was 15% of a maximal number of all neurons identified in mouse (Fig. 6, A). We found 70% GCaMP-expressing neurons two days after the recombination event. The total number of Fos-TRAPed neurons reached 15, 35 and 82 for three mice by three days after the recombination event and remained at the same level during the following imaging sessions (Fig. 6, A). Thus, the maximum level of GCaMP accumulation occurs on the third day after the Cre-recombination and persists thereafter. Based on this data, we started calcium imaging sessions in the Fos-TRAPed neurons from the fourth day after the FC-induced recombination.

**Fig. 6.**
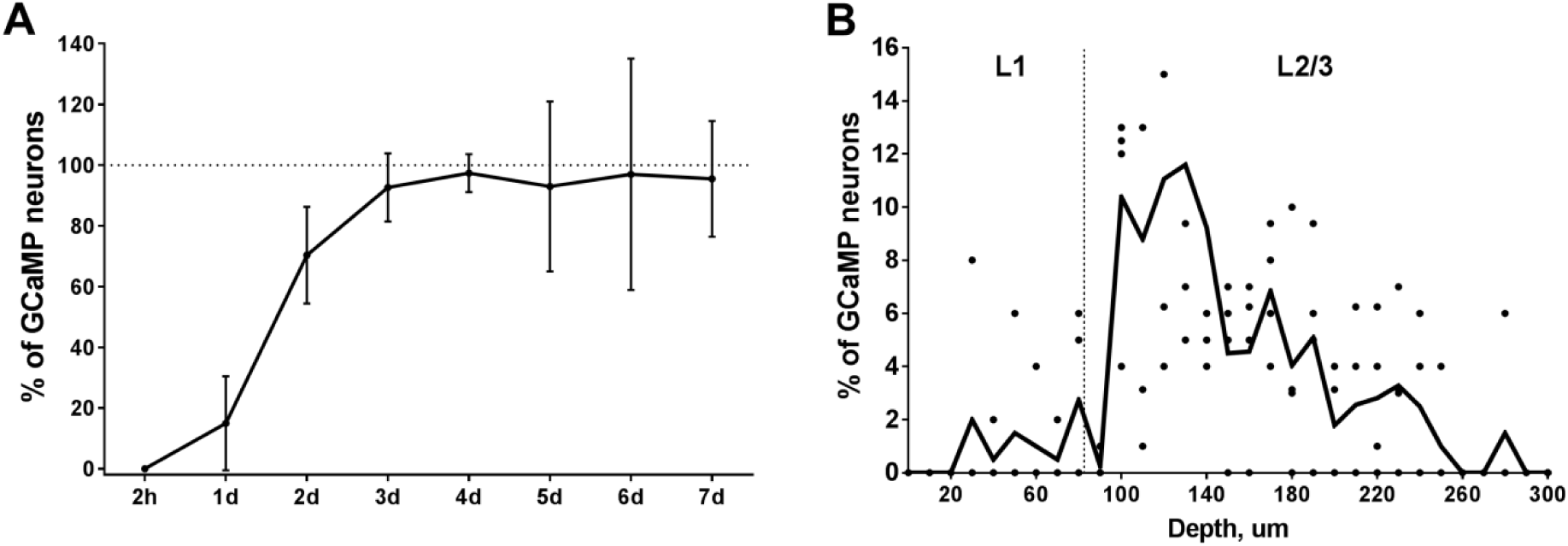
A: *GCaMP accumulation in the PtA neurons which expressed c-fos after fear conditioning. The total number of the GCaMP-positive neurons reached maximum on the fourth day after the tamoxifen injection* (Mean, 95% CI, n=3). *B:* Depth distribution of the GCaMP-expressing neurons in the *PtA (n=4).*

### Two-photon imaging of TRAPed neurons in the PtA

A stack of images was acquired in the PtA area before calcium imaging. The number of registered Fos-TRAPed neurons varied in the examined animals (20, 30, 60, 70 and 86 neurons per mouse in 0.08 mm^3^). Neurons were visible at depth up to 450 μm. The maximum density of Fos-TRAPed GCaMP-expressing neurons was detected at 100-180 μm from the brain surface, the depth that matches L2/3 of the association cortex (Fig. 7, B). Fos-TRAPed neurons showed fluorescence in the neuronal soma (nearly 15 μm in diameter), as well as in the processes (Fig. 7). The most of Fos-TRAPed neurons were pyramidal according to the observed morphology (Fig. 7). This result is consistent with the data about the types of FC-induced Fos-TRAPed neurons in the mouse neocortex (Ivashkina et al., 2018). Also, we visualized dendritic shafts (1-2 μm in diameter) with spines at depth of 5-20 μm under the brain surface (Fig. 7).

**Fig. 7.**
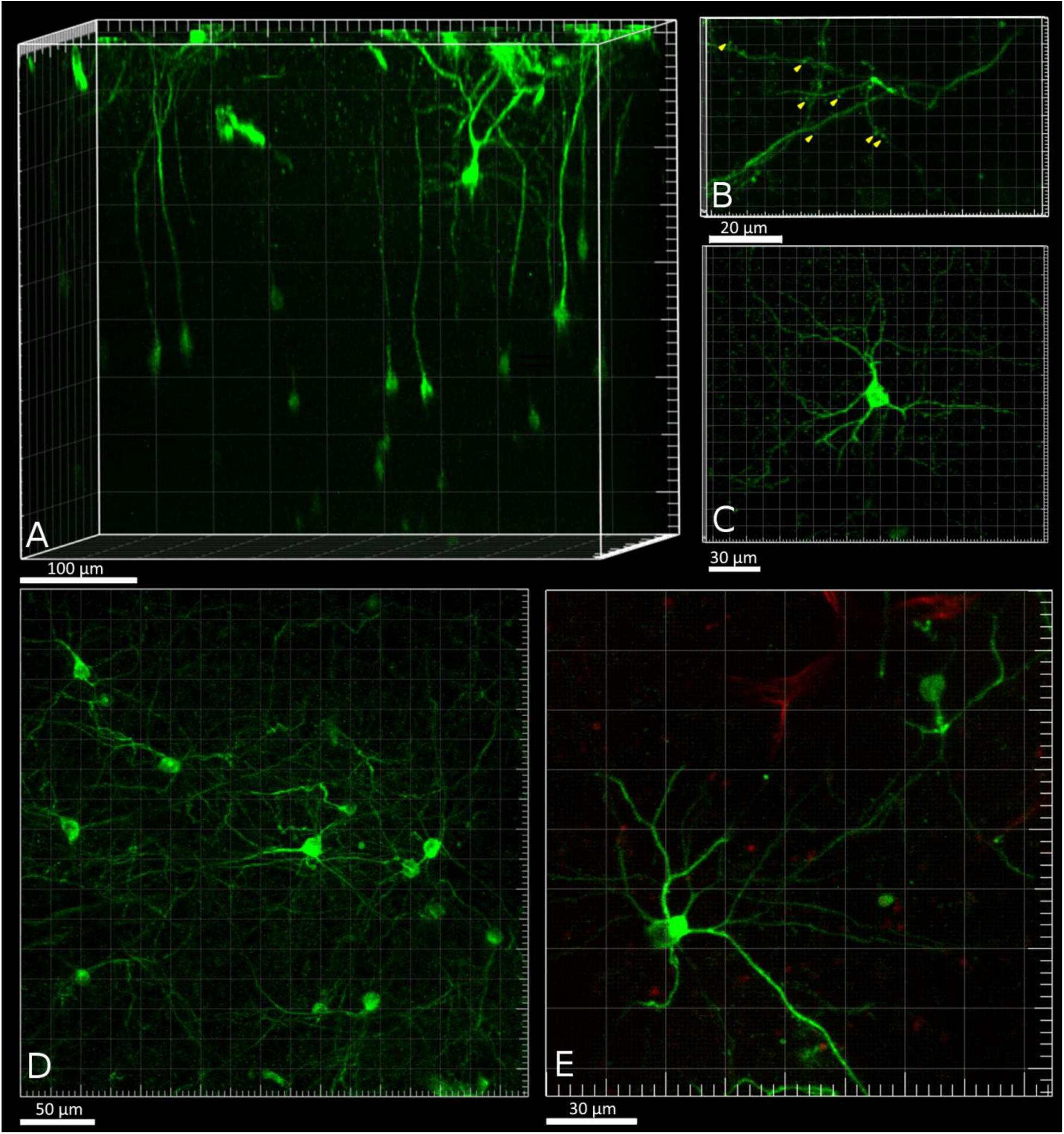
A: 3D reconstruction of GCaMP-expressing neurons in the PtA. B: Dendritic shafts and spines (tagging by yellow arrows) at L1 of the PtA. C, D, E: Examples of GCaMP-positive neurons in L2 the PtA.

To examine calcium activity in the TRAPed PtA neurons during memory recall we recorded GCaMP fluorescence during presentation of CS to head-fixed mice on the fourth day after FC. In total, we recorded calcium activity in somas of 28 neurons in L2/3 of PtA. We found 11 calcium events in six neurons (Fig. 8). However, out of these neurons no specific increase of calcium activity (i.e. increase during presentations of the CS) has been observed. Most of the neurons (75%) showed no calcium spikes during imaging sessions. However, the spontaneous calcium transients were registered in dendritic shaft and spines of the Fos-TRAPed neurons. Fig. 9 shows an increase of GCaMP fluorescence in one dendritic spine and the following increase of fluorescence in the neighboring areas of the dendrite.

**Fig. 8.**
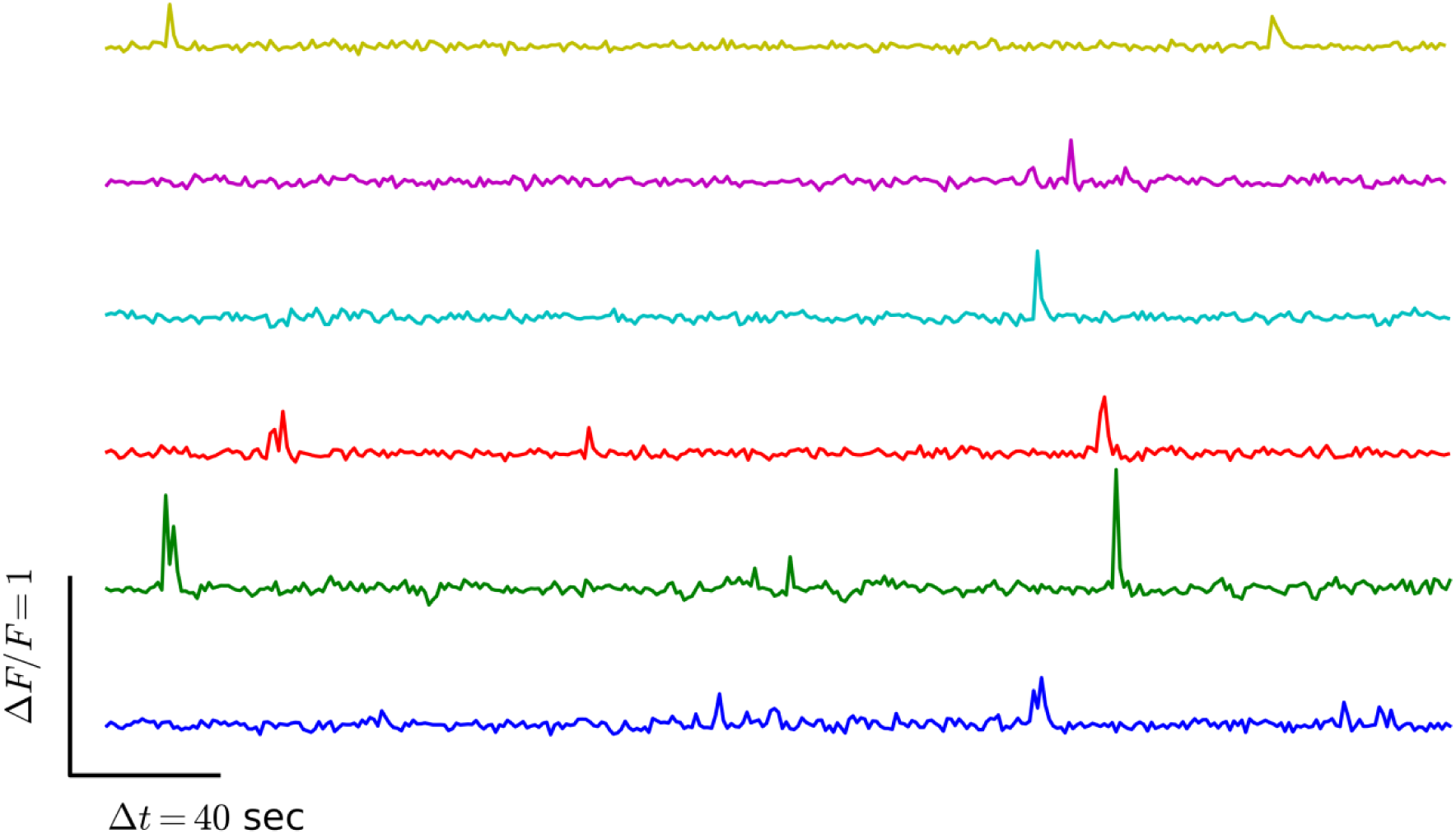
Example of calcium traces in TRAPed the PtA neurons during spontaneous activity.

**Fig. 9.**
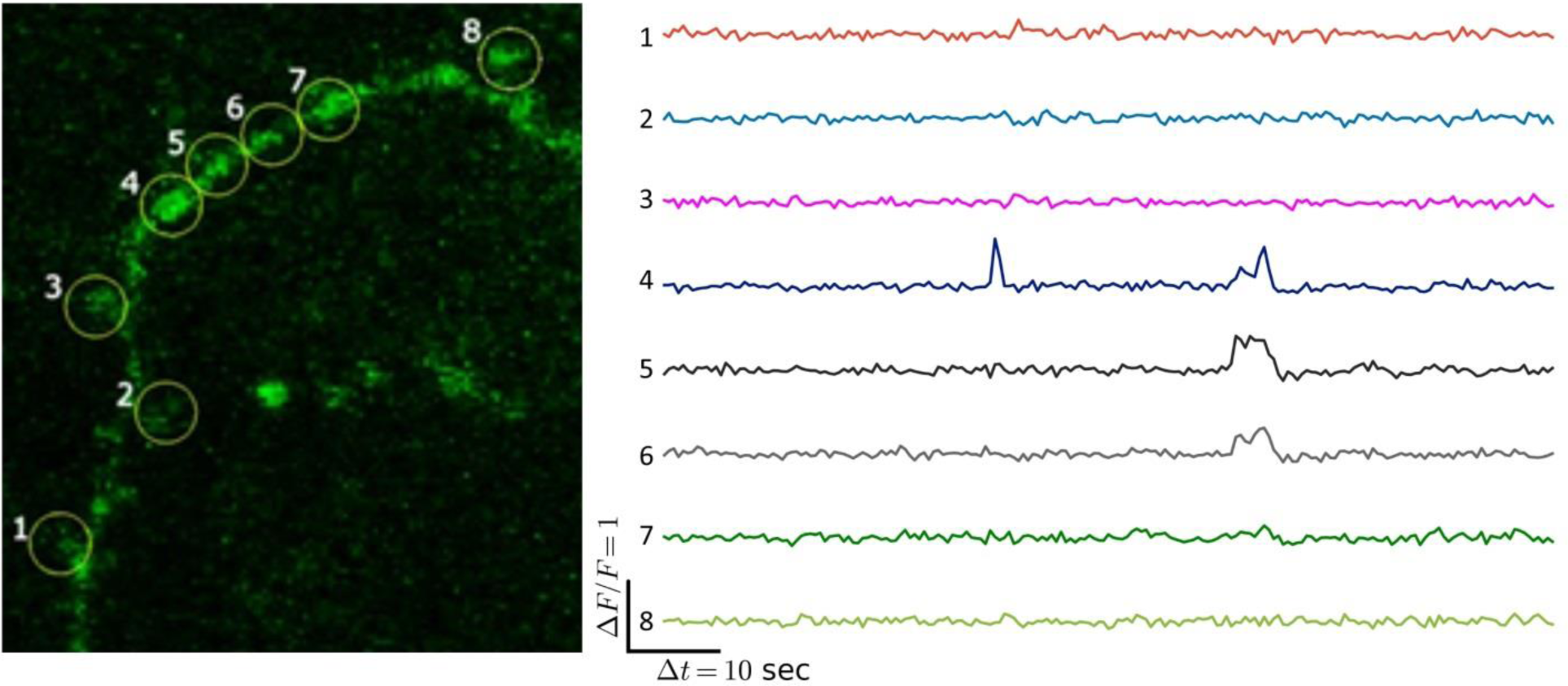
Example of calcium traces in the dendritic spines of PtA TRAPed neuron. We registered calcium event in the area 4 and following fluorescence increase in the neighboring areas 5, 6 of dendritic shaft.

Taken together, these results suggest that the Fos-Cre-GCaMP mice are suitable for investigation of calcium activity in the neurons, which were specifically activated during a particular learning episode.

## Discussion

Although it is known that rodent parietal cortex is involved in various forms of sensory processing and decision-making tasks, it is less clear how this area contributes to associative memory encoding and retrieval (Hovde et. al, 2018; Lyamzin, Benucci, 2019). Our findings suggest that the anterior part of parietal cortex – the parietal association cortex (PtA) is specifically activated during episodes of the sound cue associative fear memory encoding and retrieval. This conclusion is supported both by immunohistochemical analysis of c-Fos expression and by in-vivo activity imaging in Fos-eGFP mice.

According to previous studies the parietal cortex participates in multisensory processing and receives diverse projections from other sensory brain regions (Lyamzin, Benucci, 2019, Hovde et. al, 2018). Therefore, the increased number of Fos-expressing cells in PtA after presentation of sound stimulus during fear conditioning or in memory test could reflect this sensory processing function of the parietal cortex. To eliminate this explanation, we compared changes of the PtA activity in the Shock group with the No shock control group, which received the same procedures except the foot shock exposure.

Using Fos-immunohistochemistry we found that PtA neurons are specifically activatied in the Shock group during the memory retrieval test. In vivo Fos-eGFP imaging additionally showed specific changes of activity during both fear conditioning and memory retrieval. These results are consistent with the idea that PtA is specifically involved in encoding and retrieval of associative memory. The divergent results of the two imaging approaches can be explained by different methods used for estimation of cell populations: using ex vivo Fos immunohistochemical analysis we compared the whole populations of activated neurons, while with in vivo analysis we compared cells, which were inactive before and changed their activity specifically during fear conditioning or memory test. Noticeably, in Fos-eGFP mice we showed that the predominant proportion of the identified neurons consistently expressed Fos-eGFP during all two-photon imaging sessions. This result is consistent with the previous studies (Barth et al., 2014, Milczarek et al., 2018). To fully address this issue the estimation of the fluorescence intensity dynamics in the same neurons of PtA is needed to be done.

Previously, a laminar reorganization of neuronal activity in the mouse parietal cortex was shown due to consolidation from the recent to remote memory (Maviel et al., 2004). The shift in the pattern of neuronal activation occurred from the deep to more superficial cortical layers. Here using the Fos-immunohistochemistry imaging approach across all the cortical layers we found that the PtA was activated during recent memory retrieval. On the other hand, using in vivo Fos-eGFP imaging of superficial cortical layers we did not observe such activation. It is consistent with the idea that more superficial layers, which have a connection with other cortical areas, are less involved in the recent memory retrieval (Maviel et al., 2004, Frankland, Bontempi, 2005).

Additionally, we describe a novel transgenic mouse approach for long-term investigation of calcium activity in the neurons specifically tagged during particular cognitive episodes. Currently different IEG-based methods such as catFISH or TetTag and TRAP strategies are used to label neurons that were activated during two behavior episodes (Guzowski et al., 1999; Guenthner et al., 2013; Denny et al., 2014; Josselyn et al., 2015, Saidov et al., 2019). The catFISH has limited tagging window that allows comparing populations of neurons that were activated in two episodes with only about 30 minutes time-window (Guzowski et al., 1999). Also, catFISH is not suitable for investigation of the dynamic activity of labelled cells. Other approaches like TetTag and TRAP strategies allow comparing neuronal populations which were activated in two episodes spaced for at least 72 h (Denny et al., 2014, Guenthner et al., 2013). In this study we used TRAP strategy to perform calcium imaging of neurons, which had expressed c-Fos during the episode. We showed that these mice are suitable for imaging of calcium activity in labeled cells.

All together our results indicate the important role of the parietal association cortex in fear learning and allow us to consider the PtA as a potential target for fear- and anxiety-related psychiatric disorders.

## Acknowledgments

This work was supported by Russian Foundation for Basic Research (RFBR) grant No. 17-04-02054 A to O.I.I., RFBR grant No. 17-00-00215 to K.V.A., RSF grant No. 16-15-10323 to Fedor Subach We are grateful to Arseniy Gruzdev for help with calcium data analysis.

